# Identification of anti-resorptive GPCRs by high-content imaging in human osteoclasts

**DOI:** 10.1101/2024.12.03.626553

**Authors:** Maria L. Price, Rachael A. Wyatt, Joao Correia, Zakia Areej, Maisie Hinds, Ana Crastin, Rowan S. Hardy, Morten Frost, Caroline M. Gorvin

**Affiliations:** Department of Metabolism and Systems Science and Centre for Diabetes, Endocrinology and Metabolism (CEDAM), University of Birmingham, Birmingham, B15 2TT, UK; Centre for Membrane Proteins and Receptors (COMPARE), Universities of Birmingham and Nottingham, Birmingham, B15 2TT, UK; Molecular Endocrinology Laboratory (KMEB), Department of Endocrinology, Odense University Hospital, DK-5000, Odense C, Denmark; Department of Clinical Research, Faculty of Health Sciences, University of Southern Denmark, DK-5000, Odense C, Denmark; Department of Biomedical Sciences, University of Birmingham, Birmingham, B15 2TT, UK; National Institute for Health and Care Research (NIHR) Biomedical Research Centre (BRC), University of Birmingham, B15 2TT, UK

**Keywords:** Bone resorption, FFAR2, FFAR4, FPR1/2/3, orphan GPCRs, NFATc1 nuclear translocation, TRAP activity

## Abstract

Osteoporosis diagnoses are increasing in the ageing population and although several treatments exist, these have several disadvantages, highlighting the need to identify new drug targets. G protein-coupled receptors (GPCRs) are transmembrane proteins whose surface expression and extracellular activation make them desirable drug targets. Our previous studies have identified 144 GPCR genes to be expressed in primary human osteoclasts, which could provide novel drug targets. The development of high-throughput assays to assess osteoclast activity would improve the efficiency at which we could assess the effect of GPCR activation on human bone cells and could be utilised for future compound screening. Here we assessed the utility of a high-content imaging (HCI) assay that measured cytoplasmic-to-nuclear translocation of the nuclear factor of activated T cells-1 (NFATc1), a transcription factor that is essential for osteoclast differentiation and resorptive activity. We first demonstrated that the HCI assay detected changes in NFATc1 nuclear translocation in human primary osteoclasts using GIPR as a positive control, then developed an automated analysis platform to assess NFATc1 in nuclei in an efficient and unbiased manner. We assessed six GPCRs simultaneously and identified four receptors (FFAR2, FFAR4, FPR1, GPR35) that reduced osteoclast activity. Bone resorption assays and measurements of TRAP activity verified that activation of these GPCRs reduced osteoclast activity, and that receptor-specific antagonists prevented these effects. These studies demonstrate that HCI of NFATc1 can accurately assess osteoclast activity in human cells, reducing observer bias and increasing efficiency of target detection for future osteoclast-targeted osteoporosis therapies.

## Introduction

Osteoporosis is a common skeletal disorder characterised by reduced bone density and increased fracture risk. Over 10 million people in the United States and >25 million in Europe are estimated to have osteoporosis with >6 million fractures reported each year (Park, et al. 2023; Willers, et al. 2022). Pharmacological treatment focuses on prevention of bone loss or increases in bone mass using anti-resorptive, anabolic or combined anti-resorptive and anabolic drugs including bisphosphonates, parathyroid hormone-related analogues, and sclerostin inhibitors (Park et al. 2023). However, although effective in lowering fracture risk and increasing bone mass these drugs do have disadvantages. Bisphosphonates are associated with adverse effects including acute reactions in ∼18% of patients, and osteonecrosis of the jaw and atypical femur fractures (Khosla, et al. 2012). Denosumab is a monoclonal antibody against receptor activator of nuclear factor-κB ligand (RANKL), which inhibits osteoclast differentiation and activity (Kendler, et al. 2022). It has long-term improvements in bone mineral density; however, cessation of treatment induces rapid bone loss and increases fracture risk (Ferrari and Langdahl 2023). Romosozumab is an antibody against sclerostin which transiently inhibits bone resorption and promotes bone formation. Although more effective than other osteoporosis treatments in clinical trials, romosozumab has been associated with serious cardiovascular events. Moreover, prior treatment with bisphosphonates and denosumab attenuate its ability to increase bone mineral density (BMD) (Ebina, et al. 2024). Teriparatide increases bone mineral density, however, the efficacy is limited when used after bisphosphonates (Finkelstein, et al. 2010) and is associated with hypercalcaemia (Wen, et al. 2024). These adverse effects of current therapies indicate that additional osteoporosis therapies, which could be used alone or as part of a sequential therapy approach, would be welcomed.

G protein-coupled receptors (GPCRs) are transmembrane proteins that respond to a range of extracellular stimuli including neurotransmitters and hormones. Their cell surface expression and extracellular activation makes GPCRs inherently druggable, and GPCRs currently account for one-third of all FDA approved drugs (Hauser, et al. 2017), including teriparatide and abaloparatide that target the parathyroid hormone receptor. Our previous studies of gene expression in primary human osteoclasts at four developmental stages using RNA-sequencing revealed that 144 GPCRs are expressed in osteoclasts, including many with unknown functions in bone (Hansen, et al. 2024). Examination of the effects of three GPCRs in this study identified previously unrecognised regulators of osteoclast differentiation and activity (Hansen et al. 2024). This suggests that GPCRs that are highly expressed in bone cells could be effective targets for osteoporosis therapies and that there is a need to clarify the role of these GPCRs in bone cells.

Ideally, the assessment of multiple GPCR agonists and antagonists on osteoclast activity would be measured simultaneously and analyses would be automated. There are several methods that are commonly used to assess osteoclast activity although they differ in the extent to which they could be automated. Measurements of resorption on bone slices following incubation with osteoclasts is a routine method to assess resorption. Osteoclasts produce round pits or long trenches that can be visualised by adding toluidine blue stain (Merrild, et al. 2015; Rumpler, et al. 2013). However, bone resorption measurements are time consuming, bone slices have artefacts that are unrelated to osteoclast activity that make it difficult for untrained observers to count resorption areas accurately, as well as providing difficulties when developing automated analyses. Moreover, scanning data may be required to accurately assess the depth of resorption sites, which renders the technique unsuitable for high-throughput analyses (Rumpler et al. 2013; Sieberath, et al. 2020). Other methods to assess osteoclast differentiation and activity include quantifying tartrate-resistant acid phosphatase (TRAP) expression. TRAP is expressed by osteoclasts and has traditionally been detected using histochemistry and light microscopy. Although commonly used, quantification of TRAP-positive cells is time consuming and subject to operator bias as it can be difficult to clearly observe nuclei, which is required to designate cells as mature multinucleated osteoclasts (Cohen-Karlik, et al. 2021; Filgueira 2004). While the use of TRAP fluorescent dyes or combined treatments with nuclei stains has improved this, image quality can still be variable, and some researchers now prefer to quantify TRAP activity in cell culture media (Dai, et al. 2018; Hansen, et al. 2023). This latter method has the advantage that it can readily be adapted to high throughput assays and generates quantitative data, but it may need to be combined with other methods to demonstrate osteoclast activity (Sieberath et al. 2020). High-content imaging (HCI), which can measure the signalling effects of several treatments on the same cells on a single plate could be an alternative high-throughput method to assess osteoclast activity.

HCI allows microscopy to be performed in cell culture plates (e.g. 96-well) in a reproducible and unbiased manner. This enables the simultaneous monitoring of the effects of multiple treatments (e.g. agonists and antagonists) on cells in a high-throughput, standardised manner (Garner 2020). Signalling pathways that can be measured by HCI include monitoring calcium dynamics (Ritter, et al. 2020) and assessing nuclear translocation of signal proteins (Njikan, et al. 2018). In osteoclasts, the RANKL-mediated nuclear translocation of the nuclear factor of activated T cells-1 (NFATc1 or NFAT2) transcription factor is essential for osteoclast differentiation and resorptive activity (Ikeda, et al. 2004). We hypothesised that the quantification of NFATc1 in nuclei of osteoclasts exposed to different GPCR agonists and antagonists by HCI would enable us to screen multiple GPCRs simultaneously to assess their effects on osteoclast activity. This method would have the advantage of being able to accurately determine which cells were multinucleated by DAPI staining and assessing the activity of these osteoclasts in parallel. Here, we sought to establish an NFATc1 nuclear translocation assay in primary human osteoclasts to determine the effect of six GPCRs identified in our previous RNA-seq analyses.

## Materials and Methods

### Compounds

The working concentration and source of compounds are detailed in Table 1.

**Table 1.** Details of compounds used in these studies.

| **Agonist/**<br>**antagonist** | **Compound** | **Receptor** | **Concentration** | **Supplier and product**<br>**code** |
| --- | --- | --- | --- | --- |
| **Agonist** | 4-CMTB | Free fatty acid receptor 2<br>(FFAR2) | 10 μM | Tocris (Abingdon, UK),<br>4642/10 |
| | DL-175 | GPCR 84 (GPR84) | 10 μM | Tocris, 7082/10 |
|  | GIP | Gastric Inhibitory Polypeptide<br>Receptor (GIPR) | 10 nM | Tocris, 2084/1 |
| | N-Formyl-Met-Leu-Phe (fMLF) | Formyl peptide receptor<br>(FPR) | 1 μM | Cayman Chemical (Ann<br>Arbor, MI), CAY21495 |
| | TUG-891 | Free fatty acid receptor 4<br>(FFAR4) | 10μM | Tocris, 4601/10 |
| | Zaprinast | GPCR 35 (GPR35) | 10 μM | Merck, 37762-06-4 |
| **Antagonist** | AH7614 | Free Fatty Acid Receptor 4<br>(FFAR4) | 20 μM | Tocris, 5256/10 |
| | Boc-MLF | Formyl Peptide Receptor 1<br>(FPR1) | 5 μM | Tocris, 3730/1 |
| | GIP(3-30)NH<sub>2</sub> | Gastric Inhibitory Polypeptide<br>Receptor (GIPR) | 10 μM | CASLO (Lyngby,<br>Denmark) |
| | GLPG0974 | Free Fatty Acid Receptor 2<br>(FFAR2) | 10 μM | Tocris, 5621/10 |
| | GLPG1205 | GPCR 84 (GPR84) | 10 μM | MedChemExpress<br>(Sollentuna, Sweden),<br>HY-135303-10mg |
| | ML 145 | GPCR 35 (GPR35) | 10 μM | Tocris, 4172/10 |
| | WRW4 | Formyl Peptide Receptor 2<br>(FPR2) | 10 μM | Tocris, 2262/1 |

### Cell Culture

All cells were maintained at 37°C and 5% CO_2_. Primary human osteoclasts were differentiated from monocytes, isolated from leukocyte cones obtained from anonymous blood donations from the NHS Blood and Transplant service. Approval for isolation of monocytes from human PBMCs and their differentiation into osteoclasts was obtained from the local ethics committee in the UK (REC: 23/WA/0063, IRAS Project ID: 321094). Monocytes were enriched using the RosetteSep Human Monocyte Enrichment Cocktail (StemCell Technologies, Cambridge, UK) and separated on a Ficoll-Paque gradient (VWR, Lutterworth, UK), as described (Hansen 2025). Monocytes were seeded in α-minimal essential medium (αMEM, Gibco, Waltham, MA) supplemented with 10% newborn calf serum (NBCS, Gibco), 1% penicillin/streptomycin (P/S, ThermoFisher, Waltham, MA) and 25 ng/µL macrophage colony stimulating factor (M-CSF, BioTechne, Abingdon, UK). Monocytes were differentiated into primary human osteoclasts over 10 days, with media refreshed every 2-3 days and receptor activated nuclear factor κB ligand (RANKL, 25 ng/µL (PeproTech, London, UK) added on differentiation day 8 to stimulate osteoclast formation.

AdHEK293 cells were maintained in Dulbecco’s Modified Eagle Medium (DMEM, Gibco) supplemented with 10% NBCS. AdHEK293 cells were routinely tested to ensure they were mycoplasma-free using the TransDetect Luciferase Mycoplasma Detection kit (Generon, Slough, UK).

### Cell Viability Assays

Adherent HEK293 cells were seeded in clear-bottomed, white-walled 96-well plates, and exposed to agonists or antagonists for 72 hours at concentrations detailed in Table 1. A subset of cells were exposed to 10% DMSO as a positive control. CellTiterGlo assays (Promega, Southampton, UK) were performed according to manufacturer’s instructions and luminescence values measured on a GloMax Discover Plate Reader (Promega, Southampton, UK).

### High Content Imaging (HCI)

Mature osteoclasts (differentiation day 10) were seeded at 50,000 cells per well in clear-bottomed, black-walled 96-well plates and allowed to settle for a minimum of 4 hours. Cells were pre-incubated with vehicle or antagonist for 1 hour, followed by exposure of cells to vehicle or agonist for 1 hour. Cells were fixed with 4% PFA/PBS (Thermo Scientific/ Life Technologies, Paisley, UK), permeabilised with 0.1% Triton-X100 (Sigma-Aldrich) in PBS and blocked with 10% donkey serum in PBS. Primary incubation with a rabbit polyclonal anti-NFATc1 antibody (1:120, ab25916, Abcam, Cambridge, UK) was performed for 1 hour, then with a donkey anti-rabbit secondary AlexaFluor488 for 1 hour (1:500, A21206, Invitrogen, Waltham, MA). Nuclei were stained with NucBlu™ DAPI for 15 minutes (1:100, Invitrogen; R37606). Cells were imaged on a Cell Discoverer 7 (Zeiss, Oberkochen, Germany), using a Axiocam 702 camera, Plan-Apochromat objective (at 40x magnification), and LED light illuminators to measure DAPI (excitation 353nm, emission 465nm) and AlexaFluor488 (excitation 493nm, emission 517nm). Images were captured using Zen Black software (Zeiss) programmed to take five images of each well. Two technical repeats were performed for each treatment and the location of each treatment on the plate varied between biological repeats.

Manual analysis was performed of wells exposed to glucose-dependent insulinotropic polypeptide (GIP) or the GIP-receptor (GIPR) antagonist using ImageJ (NIH) to measure total corrected cellular fluorescence (TCCF), accounting for cell area and background signal, as previously described (McCloy, et al. 2014). In brief, ImageJ was used to draw an outline around each cell and around an area of background, mean fluorescence was measured in both regions of interest (ROIs), then the total corrected cellular fluorescence (TCCF) was calculated as: fluorescent signal in the ROI – (area of selected cell x mean fluorescence of background readings). This was then repeated to measure the area around each nucleus. Nuclear values were subtracted from cytoplasmic values to derive a TCCF for the nucleus and the cytoplasm. Nuclear/ cytoplasmic ratios were then derived and plotted in GraphPad Prism.

Automated analysis to measure signal inside nuclei was performed using Zeiss arivis Pro Software. Fluorescent channels were separated and software was used to identify the nuclei as regions of interest using the DAPI channel. The signal intensity was then measured in the NFAT-488 channel. The analysis platform for quantifying nuclear fluorescence using arivis is available at the open science framework https://osf.io/43a8h/. Only nuclei from mature multinucleated osteoclasts were used for data analysis.

### Osteoclast Resorption Assays

Mature osteoclasts (differentiation day 10) were seeded at a density of 50,000 cells per well on bovine cortical bone slices (Boneslices.com, Jelling, Denmark) in 96-well plates and allowed to settle for 1 hour. GPCR agonists and/or antagonists were added and plates were incubated for 72 hours at 37 °C and 5% CO_2_. Media was removed and dH_2_O added to terminate the experiment, then cells removed from the bone slices with a cotton swab. Bone slices were stained with toluidine blue solution (1% toluidine blue, 1% sodium borate in dH_2_O, both from Sigma-Aldrich, Gillingham, UK) for 20 seconds. Bone slices were visualised on an Olympus BX53 microscope (Olympus, Tokyo, Japan) at 10x magnification and areas of bone resorption were quantified using a 10×10 counting grid (24.5 mm, Graticules Optics Ltd., Tonbridge, UK). Measurements were made in eight predefined areas spanning the bone slice, with the observer blinded to the treatment conditions, and resorption was expressed as the percentage eroded surface per bone slice.

### Tartrate-resistant acid phosphatase 5b (TRAP) activity assays

Measures of TRAP activity were performed as described (Hansen et al. 2024). Mature osteoclasts (differentiation day 10) were seeded at a density of 50,000 cells per well in 96-well plates and exposed to agonists and/or antagonists for 72 hours. Conditioned media was collected and 10 µL transferred to a clear 96-well plate, then 90 µL TRAP solution buffer (1M acetate, 0.5% Triton X-100, 1M NaCl, 10 mM EDTA, 50 mM L-Ascorbic acid, 0.2 M disodium tartrate, 82 mM 4-nitrolphenylphosphate, all Sigma-Aldrich) added prior to incubation in the dark for 30 minutes at 37 °C. Reactions were stopped with 0.3M NaOH (Sigma-Aldrich) and TRAP activity measured at absorbance 400 nm and 645 nm on a SpectraMax ABS (Molecular Devices, San Jose, CA). TRAP activity was expressed as a 400/645 absorbance ratio.

### Statistical Analysis

The number of experimental replicates denoted by n is indicated in figure legends. Statistical analyses were performed using GraphPad Prism 9, with details described in figure legends. Normality tests (Shapiro-Wilk or D’Agostino-Pearson) were performed on all datasets to determine whether parametric or non-parametric statistical tests were appropriate. A p-value of <0.05 was considered statistically significant.

## Results

### Identification of GPCR targets and assessment of compound toxicity

We have previously shown that 144 GPCRs are expressed in primary human osteoclasts and that many of these are differentially expressed during osteoclast differentiation (Hansen et al. 2024). Six GPCRs were selected for analyses based on: a ready availability of agonists and antagonists, an incomplete understanding or unknown function in human osteoclasts, or because the receptors are current drug development targets and compounds could be rapidly repurposed for future osteoporosis studies. GIPR was used as a positive control in each experiment as it has previously been shown to reduce NFATc1 nuclear translocation, osteoclast resorption and TRAP activity (Hansen et al. 2023). In the RNA-seq datasets one GPCR, free-fatty acid receptor-4 (FFAR4), had an increase in gene expression during differentiation; while five receptors, FFAR2, formyl peptide receptor 1 and 2 (FPR1, FPR2), GPR35 and GPR84 had a reduction in gene expression during differentiation. We have previously shown that FFAR4 can signal in primary human osteoclasts (Hansen et al. 2024) and FFAR2 knockout mice (*Gpr43^-/-^*) have suppressed bone resorption (Montalvany-Antonucci, et al. 2019). All three members of the FPR family are expressed in primary human osteoclasts and are known to respond to the N-Formyl-Met-Leu-Phe (fMLF) peptide in other cell types (Migeotte, et al. 2006). Previous studies have indicated that formyl peptide may affect bone resorption (Park, et al. 2017), but the receptors involved in human cells remain to be clarified. GPR35 and GPR84 are defined as orphan GPCRs as their native endogenous ligands are unknown. Several orphan receptors are expressed in human osteoclasts, and GPR35 and GPR84 may negatively regulate osteoclastogenesis in mice (Park, et al. 2018). For these reasons, we chose these six receptors to screen using HCI.

To ensure that any effects observed in cell studies were due to receptor specific activity rather than a toxic effect of compounds, cell viability was first assessed in HEK293 cells. Cells were exposed to compounds for 72 hours, and cell viability measured using CellTiterGlo assays. When used alone, no agonists or antagonists significantly affected cell viability (Figure 1A-B). Treatment of cells with 10% DMSO for 72 hours, used as a positive control, did significantly reduce cell viability (Figure 1A-B). A subset of cells was also exposed to antagonists for both FPR1 and FPR2 as an alternative to suppress FPR3 activity as no robust antagonists exist against this receptor. However, combined treatment reduced the viability of the primary osteoclasts. Therefore, combined treatments were not pursued in subsequent studies.

**Figure 1.**
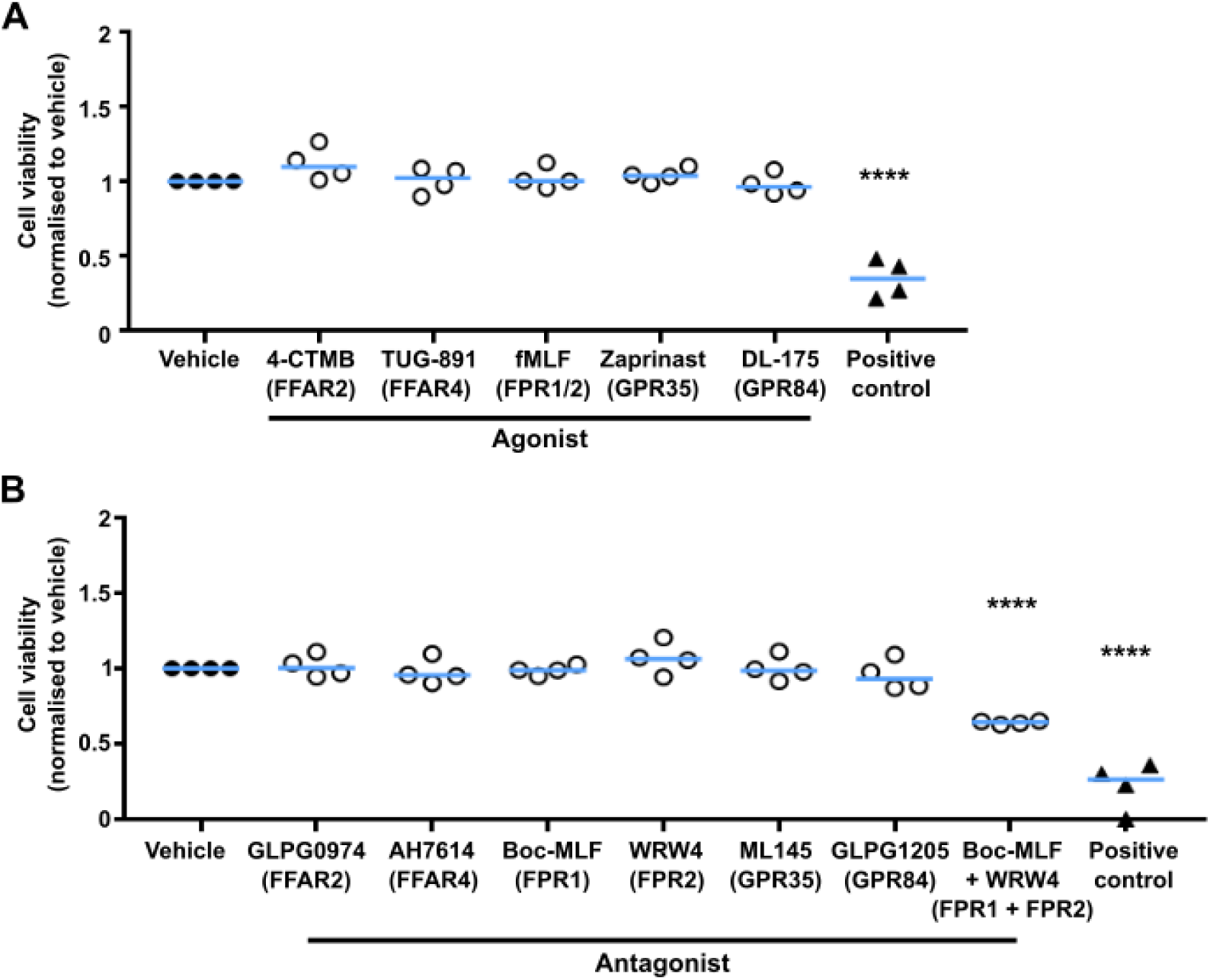
GPCR agonists and antagonists do not affect osteoclast viability Cell viability of HEK293 cells exposed to (**A**) agonists and (**B**) antagonists for the GPCRs shown in parentheses measured by CellTiterGlo. A positive control, 10% DMSO, was used, whichis known to reduce cell viability. Boc-MLF and WRW4 were combined to assess the feasibility of antagonising both FPR1 and FPR2 in combination. Cells were exposed to compounds for 72 hours and each biological replicate is shown as a separate point. Statistical analyses compared to vehicle were performed by one-way ANOVA with Dunnett’s multiple comparisons test. ****p<0.0001.

### Optimisation of the HCI technique

Translocation of NFATc1 to the nucleus to induce osteoclast-mediated gene transcription is essential for RANKL-mediated effects on osteoclast activity and differentiation (Takayanagi, et al. 2002), and GPCR agonists, including GIP, have been shown to affect the NFATc1 signalling pathway (Hansen et al. 2023). We hypothesised that measuring agonist-induced effects on NFATc1 nuclear translocation by HCI would be an efficient way to test which receptors likely affect osteoclast activity and could be adapted in future studies to screen compound libraries to identify anti-resorptive targets. To validate the HCI protocol we first assessed whether HCI could replicate our previous findings that activation of GIPR induces NFATc1 nuclear translocation (Hansen et al. 2023). Mature osteoclasts in 96-well plates were pre-exposed to the GIPR antagonist, GIP(3-30)NH_2_, then GIPR was stimulated with GIP to induce NFATc1 nuclear translocation. Fixed cells were then exposed to NFATc1 antibody and fluorescently labelled with AlexaFluor488, and counter-stained with DAPI to label nuclei prior to performing HCI (Figure 2A-B). Images were first analysed manually using a previously described technique (McCloy et al. 2014) in which fluorescence is measured in the cell cytoplasm and nucleus, and a nuclear-to-cytoplasmic ratio is derived. Consistent with previous studies, GIP was shown to reduce NFATc1 nuclear translocation (Figure 2A-D). Pre-exposure of osteoclasts to the GIPR antagonist prevented the GIP-induced reduction in NFATc1 nuclear translocation (Figure 2A-D). Therefore, the HCI replicates the data previously acquired on coverslips by confocal microscopy (Hansen et al. 2023).

**Figure 2.**
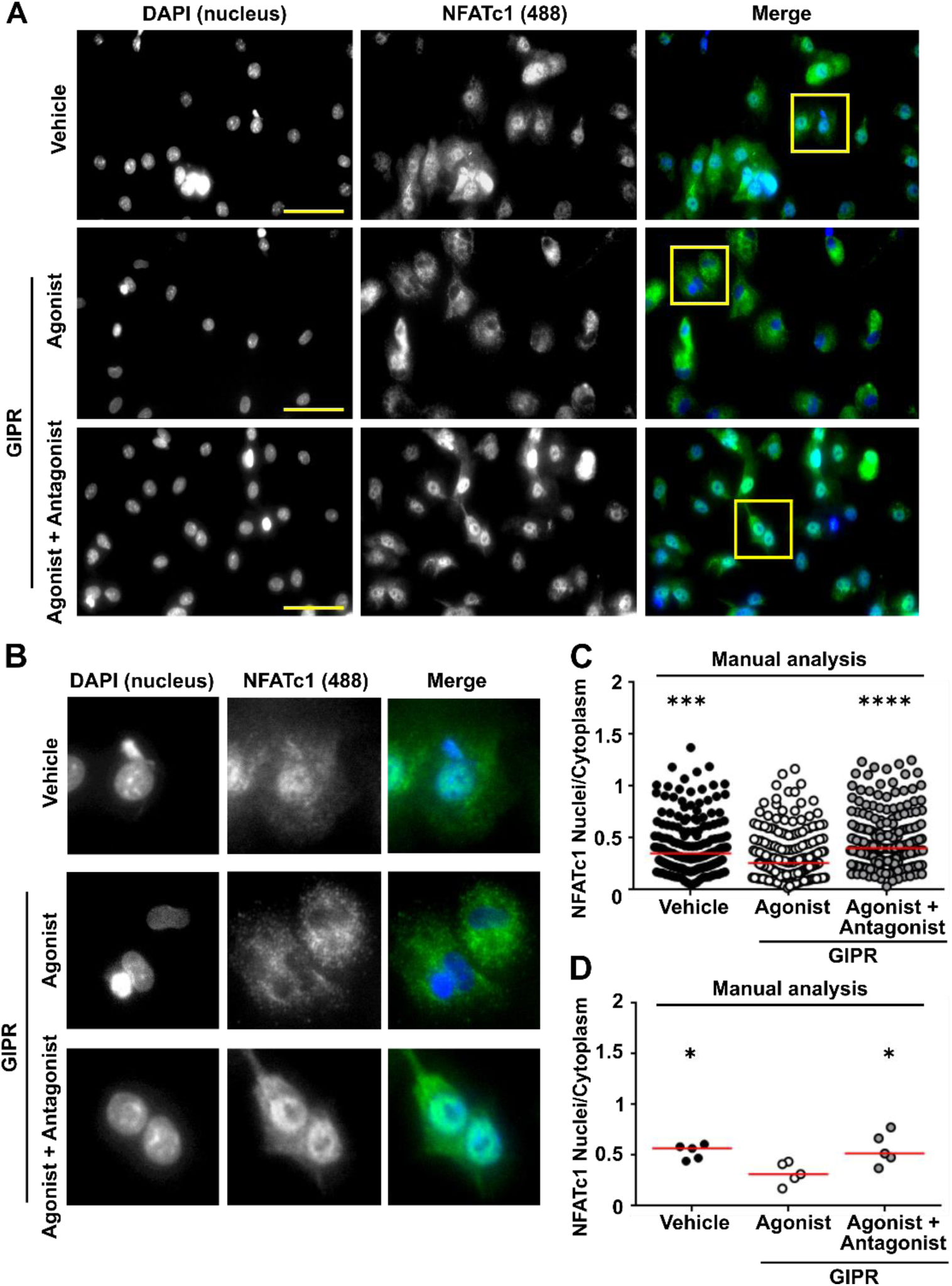
Validation of the HCI and automated analysis to detect changes in NFATc1 nuclear translocation (**A**) High content images of mature osteoclasts exposed to vehicle, GIP (agonist) or GIP with GIP(3-30)NH_2_ (antagonist), then labelled with NFATc1 and AlexaFluor488. Scale, 50 µm. (**B)** Close-up images showing NFATc1 nuclear fluorescence is reduced in cells exposed to GIP. (**C**) Manual quantification of NFATc1 in osteoclast nuclei and the cytoplasm. Vehicle (182 cells), GIP (196 cells), GIP(3-30)NH_2_ (211 cells) from 5 donors. Data shows nuclear/cytoplasmic ratios for all cells measured with median shown in red. (**D**) Quantification of nuclear/ cytoplasmic NFATc1 in individual donors. Median is shown in red. Statistical analyses were performed by Kruskal–Wallis one-way ANOVA with Dunn’s multiple comparisons test and compare responses to agonist. ****p<0.001, ***p<0.001, *p<0.05.

We then sought to automate the analysis of the high-content images to remove observer bias and increase analysis efficiency. An automated analysis pipeline was designed using Zeiss arivis Pro Software in which nuclei were identified using the DAPI images, then signal intensity was measured in the corresponding areas on the NFAT-488 image. To determine whether measuring nuclei only was a valid way to measure NFATc1 nuclear translocation, we returned to our original dataset measured by manual analysis and quantified nuclei fluorescence only. This similarly demonstrated a significant reduction in fluorescence intensity compared to cells exposed to vehicle or cells pre-treated with the GIPR antagonist (Figure 3A-B). We then tested our automated nuclei analysis platform to determine whether it could accurately detect differences between fluorescence intensities in our HCI dataset. The automated analysis detected a similar number of cells to the manual analysis and there was less variability in the data obtained. In the automated analyses GIP still had a significant reduction in NFATc1 nuclear fluorescence compared to osteoclasts exposed to vehicle or antagonist (Figure 3C-D). Thus, the automated analysis can accurately detect differences between treatments, and we proceeded to assess other GPCR agonists by HCI with automated analysis.

**Figure 3.**
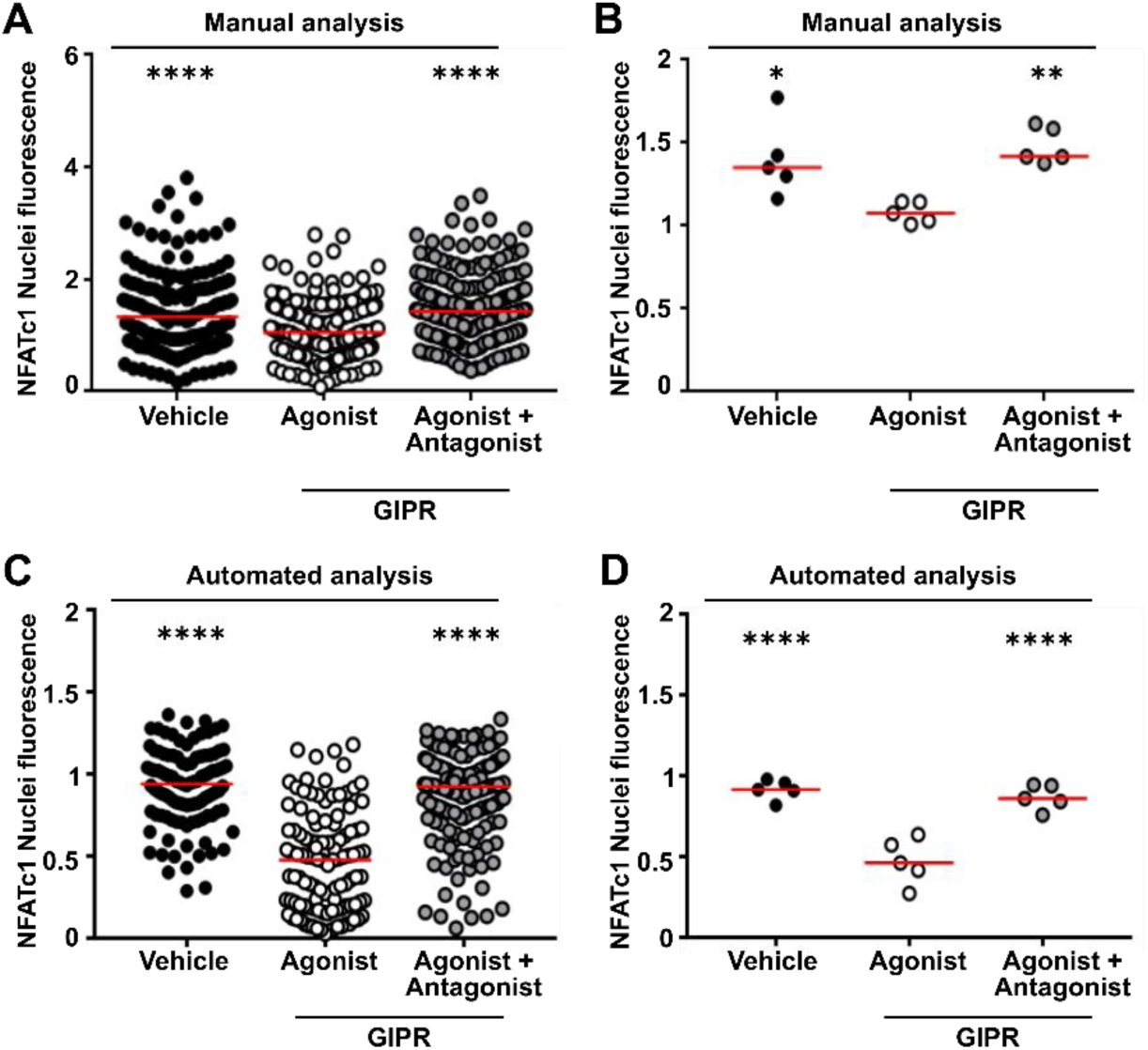
Optimisation of HCI workflow (**A-B**) Manual quantification of NFATc1 in osteoclast nuclei showing (A) all nuclei counted and (B) average for each donor. Vehicle (182 cells), GIP (196 cells), GIP(3-30)NH_2_ (211 cells) from 5 donors. (**C-D**) Quantification of NFATc1 in osteoclast nuclei using the arivis automated workflow. Vehicle (180 cells), GIP (193 cells), GIP(3-30)NH_2_ (207 cells) from 5 donors. Median is shown in red. Statistical analyses were performed by Kruskal–Wallis one-way ANOVA with Dunn’s multiple comparisons test and compare responses to agonist. ****p<0.001, **p<0.01, *p<0.05.

### High content imaging detects four GPCRs that reduce NFATc1 signalling

NFATc1 nuclear translocation, with agonists and antagonists for the six GPCRs selected from the RNA-seq dataset, was then investigated by HCI. Exposure of mature osteoclasts to agonists for FFAR2 and FFAR4 (4-CMTB and TUG-891, respectively) significantly reduced NFATc1 nuclear translocation (Figure 4A-D). Pre-incubation with receptor-specific antagonists (GLPG0974 and AH7614, respectively) prevented this reduction. Similarly, exposure of osteoclasts to fMLF, which activates all three FPRs, significantly reduced NFATc1 nuclear translocation. Pre-incubation with the FPR1 antagonist (Boc-MLF) prevented this reduction in translocation, whereas the FPR2 antagonist (WRW4) had no effect on fMLF-induced responses (Figure 4E-F). Activation of GPR35 with agonist (Zaprinast) reduced NFATc1 nuclear translocation, which was prevented by combined exposure of receptor-specific agonist and antagonist (ML 145) (Figure 4G-H). In contrast, no significant differences were observed in osteoclasts exposed to agonist (DL-175) and antagonist (GLPG1205) of GPR84 (Figure 4I-J). Therefore, HCI identified that four GPCRs reduce NFATc1 signalling in osteoclasts.

**Figure 4.**
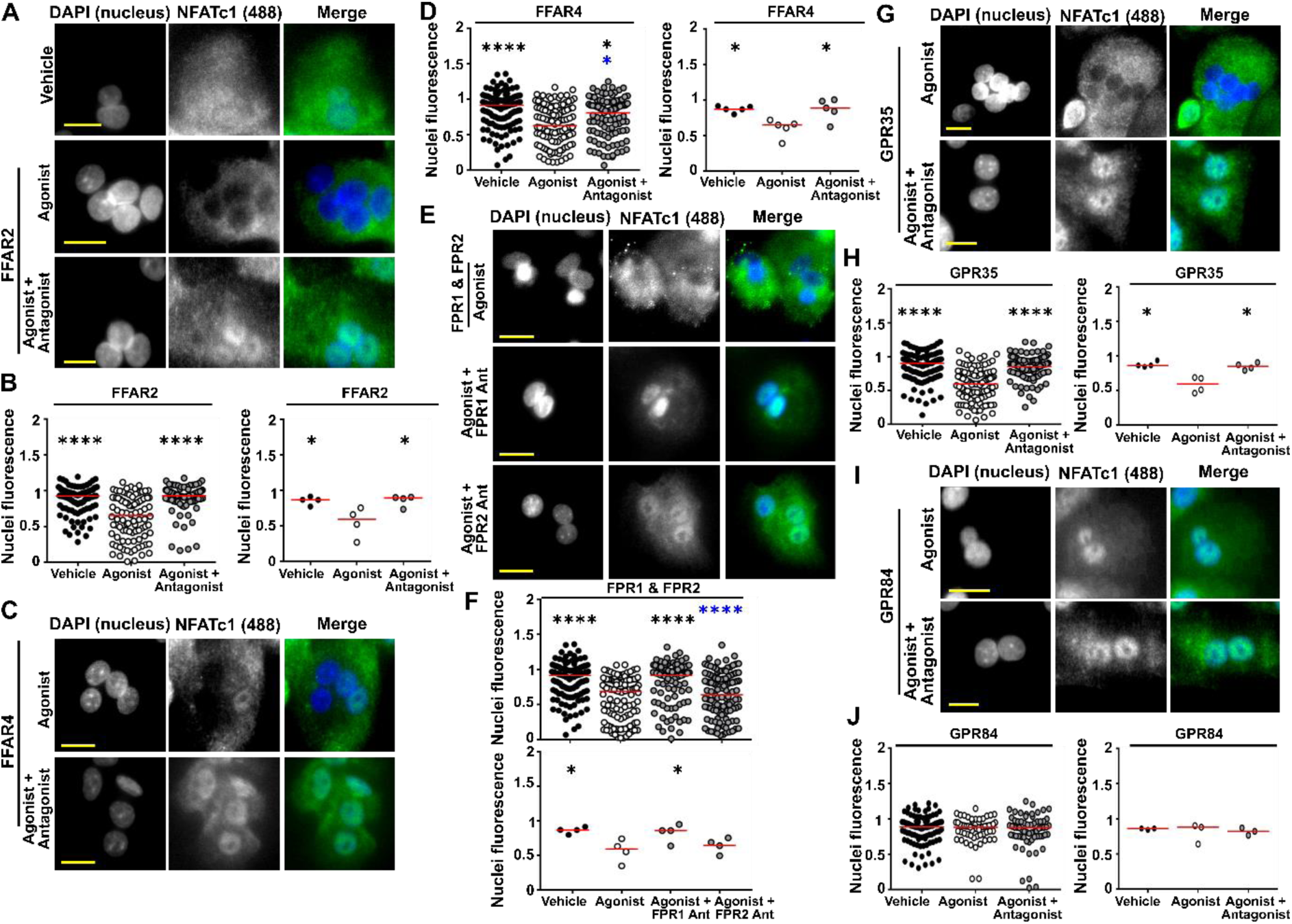
HCI identifies four GPCRs that reduce NFATc1 nuclear translocation (**A**) Images showing NFATc1 expression in mature osteoclasts exposed to vehicle and FFAR2 agonist and agonist with antagonist. (**B**) Quantification of NFATc1 in osteoclast nuclei analysed using the automated workflow showing all cells analysed (left) and average data for each donor (right). Vehicle (92 cells), agonist (94 cells), agonist with antagonist (79 cells). (**C**) Images showing NFATc1 expression in mature osteoclasts exposed to vehicle and FFAR4 agonist and agonist with antagonist. (**D**) Automated quantification of NFATc1 in osteoclast nuclei showing all nuclei analysed (left) and average data for each donor (right). Vehicle (131 cells), agonist (106 cells), agonist with antagonist (103 cells). (**E**) Images showing NFATc1 expression in mature osteoclasts exposed to vehicle and FPR1/2 agonist and agonist with antagonist. (**F**) Automated quantification of NFATc1 in osteoclast nuclei showing all cells analysed (top) and average data for each donor (below). Vehicle (131 cells), agonist (115 cells), agonist with FPR1 antagonist (ant) (95 cells) or agonist with FPR2 antagonist (111 cells). (**G**) Images showing NFATc1 expression in mature osteoclasts exposed to vehicle and GPR35 agonist and agonist with antagonist. (**H**) Automated quantification of NFATc1 in osteoclast nuclei showing all cells analysed (left) and average data for each donor (right). Vehicle (115 cells), agonist (102 cells), agonist with antagonist (106 cells). (**I**) Images showing NFATc1 expression in mature osteoclasts exposed to vehicle and GPR84 agonist and agonist with antagonist. (**J**) Automated quantification of NFATc1 in osteoclast nuclei showing all cells analysed (left) and average data for each donor (right). Vehicle (102 cells), agonist (59 cells), agonist with antagonist (64 cells). Median is shown in red. Statistical analyses were performed by Kruskal–Wallis one-way ANOVA with Dunn’s multiple comparisons test. Black asterisks show comparisons to agonist and blue to vehicle. ****p<0.001, ***p<0.001, *p<0.05. Scale, 50 µm for all images.

### The four GPCRs identified by HCI reduce osteoclast activity

The HCI studies showed that activation of four receptors (FFAR2, FFAR4, FPR1, GPR35) significantly reduced NFATc1 nuclear translocation, and may affect osteoclast resorption. To investigate whether results from HCI correlate with changes in osteoclast activity, two assays that are routinely used in osteoclast research, resorption assays on bone slices and detection of TRAP activity, were performed.

To determine whether the six selected GPCRs affected osteoclast activity, mature primary human osteoclasts were incubated with either vehicle, agonist, or agonist with antagonist for 72 hours, then toluidine blue staining and quantification of osteoclast resorption sites performed. Consistent with our previous studies, bone slices incubated with the GIPR agonist, GIP, had significantly fewer osteoclast resorption sites than bone slices exposed to vehicle or the GIPR antagonist, GIP(3-39)NH_2_ (Figure 5A-B). Incubation of cells with the FFAR2 and FFAR4 agonists also reduced resorption, and this was reversed by exposure of cells to receptor-specific antagonists (Figure 5C-D). Incubation of osteoclasts with fMLF significantly reduced bone resorption (Figure 5C-D). Incubation of cells with the FPR1-specific antagonist impaired the ability of fMLF to reduce bone resorption, but there was a significant difference compared to vehicle-treated cells, indicating another FPR may be involved. The FPR2-specific antagonist had no effect, suggesting that FPR3 may have a role in suppression of osteoclast resorption. Activation of GPR35 and GPR84 also reduced human osteoclast resorption (Figure 5C-D). However, co-treatment of cells with receptor-specific antagonists only prevented the effect of GPR35 stimulation, but had no effect on GPR84 activation (Figure 5C-D).

**Figure 5.**
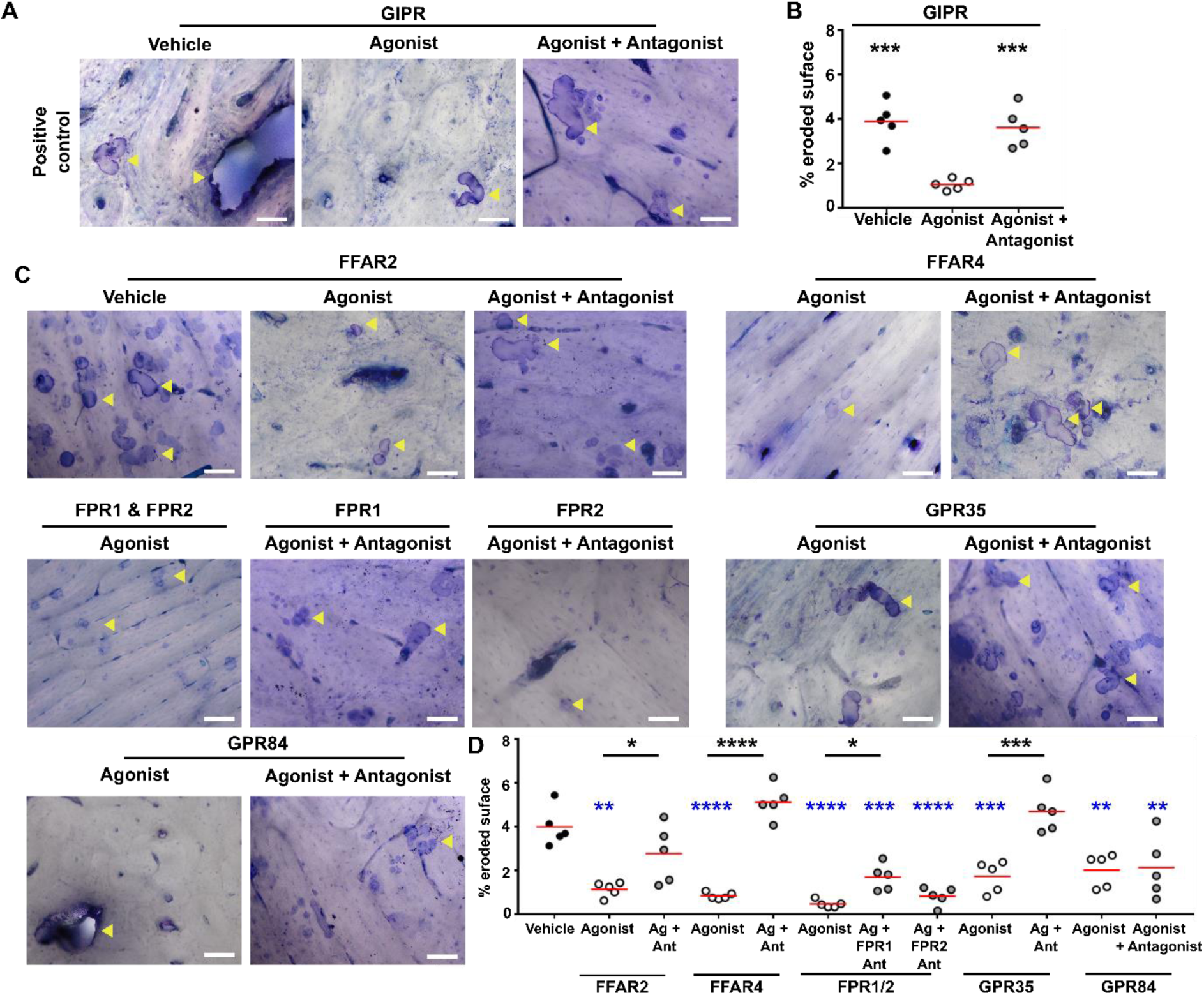
Activation of GPCRs in primary human osteoclasts reduces osteoclastic bone resorption (**A**) Representative images of resorption areas (indicated with yellow arrows) on bone slices incubated with mature osteoclasts for 72 hours and exposed to vehicle, agonist or agonist with antagonist for GIPR. (**B**) Quantification of the resorption areas on bone slices expressed as percentage eroded surface. Each point represents a single donor. (**C**) Representative images of resorption areas (indicated with yellow arrows) on bone slices incubated with mature osteoclasts for 72 hours and exposed to vehicle, agonist or agonist with antagonist for the six GPCRs, with (**D**) quantification of the percentage eroded surface. Each point represents a single donor. Median is shown in red. Statistical analyses were performed by one-way ANOVA with Tukey’s or Dunnett’s multiple comparisons test. Black asterisks show comparisons to agonist and blue asterisks to vehicle. Scale, 100 µm.

Finally, measurements of TRAP in media from osteoclasts incubated with GPCR agonists and antagonists for 72-hours was assessed. Exposure of cells to GIP was shown to reduce TRAP activity, as previously described (Hansen et al. 2023) (Figure 6A). TRAP activity was also reduced by activation with all five agonists. However, pre-exposure of cells to GPCR-specific antagonists prevented the effect of FFAR2, FFAR4, FPR1 and GPR35 on TRAP activity (Figure 6B-F). Therefore, bone resorption assays and TRAP activity assays correlate well with results from the HCI studies.

**Figure 6.**
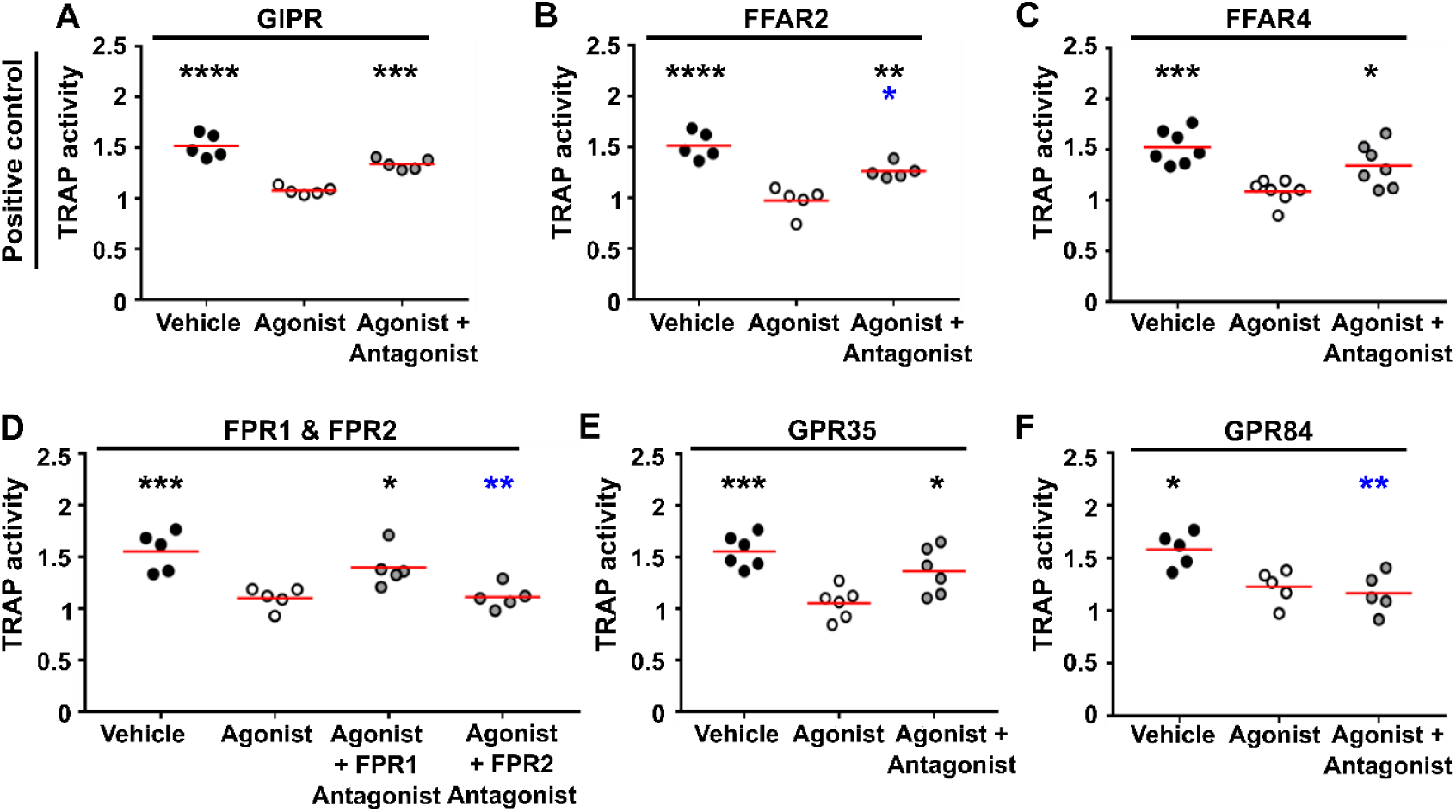
Activation of four GPCRs reduces TRAP activity Quantification of TRAP activity from mature osteoclasts exposed for 72 hours to vehicle, agonist or agonist and antagonist specifically targeting (**A**) GIPR, (**B**) FFAR2, (**C**) FFAR4, (**D**) FPR1 and FPR2, (**E**) GPR35, (**F**) GPR84. Each point represents a single donor. Median is shown in red. Statistical analyses were performed by one-way ANOVA with Tukey’s or Dunnett’s multiple comparisons test. Black asterisks show comparisons to agonist and blue asterisks to vehicle.

## Discussion

Our studies have demonstrated the utility of a HCI platform to identify regulators of osteoclast activity. Assessment of agonists and antagonists to seven GPCRs identified five receptors (GIPR, FFAR2, FFAR4, FPR1, GPR35) reduce NFATc1 activity, and these same receptors were identified to reduce bone resorption using traditional measures of osteoclast activity. Additionally, measurement of TRAP activity in cell media from osteoclasts exposed to agonists for 72 hours was as proficient as toluidine blue stained bone resorption assays at identifying GPCRs that affect osteoclast activity. Therefore, a combined assessment of the effect of compounds on NFATc1 nuclear translocation by HCI and TRAP activity in cell media would allow both the acute (one hour) and long-term (72 hour) effects of compounds on GPCRs to be assessed and should provide sufficient information to decide which receptors to assess in further detail. Moreover, this assay could be used in safety tests to identify compounds that do not affect osteoclast activity, and therefore may have fewer potential off-target effects in bone. HCI has the advantage that there is no requirement for the genetic manipulation of receptors that could have off-target effects on cell health or differentiation, these assays are not subject to observer bias and they can be performed in 96-or 384-well plates allowing multiple receptors to be assessed simultaneously. The HCI of additional proteins that undergo nuclear translocation in osteoclasts (e.g. NFκB) could be performed in parallel with NFATc1 to provide details on multiple signalling pathways within the same assay.

The assays we developed are currently semi-automated as an operator is still required to determine which cells are multinucleated (and therefore mature osteoclasts) after fluorescence in all nuclei has been obtained. While this may reduce the efficiency of data analysis, this method is still more efficient than bone resorption assays that require manual counting over many hours, whereas HCI can obtain data from 96-well plates within minutes using the latest technologies. Additionally, with the rapid development of machine learning it is possible that our automated analysis could be improved to determine not just arbitrary scoring of multinucleated vs. single-nuclei cells, but could perform sophisticated analyses determining osteoclast activity in cells with different numbers of nuclei (Cohen-Karlik et al. 2021).

Two receptors known to bind free-fatty acids were examined in these studies. We have previously shown that activation of FFAR4 reduces human osteoclast activity, while others have shown effects in mouse cell-lines (Hansen et al. 2024; Kasonga, et al. 2019). Here, we provided further insights by including an FFAR4-specific antagonist that we showed prevents TUG-891 affects on osteoclast resorption. Moreover, we demonstrated that acute activation of FFAR4 reduces NFATc1-mediated nuclear translocation, consistent with previous studies that showed FFAR4 activation attenuates NFATc1 mRNA induction (Kim, et al. 2016). FFAR2 binds short-chain free-fatty acids and has been shown in a single study to affect bone cell activity. A global *Gpr43^-/-^* mouse had a reduced number of osteoblasts, increased osteoclasts and changes in bone turnover markers (Montalvany-Antonucci et al. 2019). Short-chain fatty acids and FFAR2-specific agonists reduced osteoclast differentiation in cells isolated from wild-type mice indicating that there may be a direct effect of activation of FFAR2 on osteoclasts, but effects on human cells were not investigated, FFAR2-specific antagonists were not examined and studies were performed in only two cell cultures (Montalvany-Antonucci et al. 2019). Our studies verified these previous findings and demonstrated that human FFAR2 may regulate osteoclast activity.

We were unable to demonstrate a role for GPR84 in human osteoclasts. GPR84 is highly expressed on immune cells and may have a role in phagocytosis (Luscombe, et al. 2020). Several studies suggest GPR84 may bind medium-chain fatty acids. However, the receptor binds these ligands with low potency and is not expressed at high concentrations in many physiologically relevant tissues (Luscombe et al. 2020), therefore GPR84 is still regarded as an orphan receptor. Very few studies have investigated GPR84 in osteoclasts. One study examined osteoclasts differentiated from bone marrow-derived macrophages and showed that GPR84 overexpression suppressed osteoclast differentiation, while knockdown enhanced differentiation (Park et al. 2018). Findings from this study may differ from ours as we only used receptor-specific agonists and antagonists rather than modifying gene expression. We chose this approach as gene manipulation could affect receptor functions in precursor cells (e.g. macrophages) rendering it difficult to determine cell-specific responses, and we hypothesised that any compounds developed as osteoporosis therapies would likely target receptor activity (e.g. antagonists or allosteric modulators) rather than suppressing gene expression.

Activation of GPR35, an orphan GPCR, significantly reduced NFATc1 nuclear translocation, resorption and TRAP activity in these studies. Previous studies have indicated that *GPR35* is downregulated in individuals with osteoporosis and in mouse models of the disease (Zhang, et al. 2021) and that activation of GPR35 improves bone density in osteoporotic mice (Ma, et al. 2023). This has been attributed to the effect of GPR35 on osteoblasts as *Gpr35^-/-^*mice have reduced bone mass due to impaired osteoblast development (Zhang et al. 2021). However, osteoclast-specific effects were not examined in these studies, and our data support a role for GPR35 stimulation in also affecting osteoclast bone resorption. One disadvantage of our studies is that we used an agonist, Zaprinast, that may have off-target effects on phosphodiesterase-5 and −6 (PDE5/ 6) (Taniguchi, et al. 2006). However, PDE5 is not expressed and PDE6 is expressed at very low concentrations in mature osteoclasts (Hansen et al. 2024), and our use of a GPR35-specific antagonist increases confidence that our findings are mediated by GPR35 activity.

Our studies showed fMLF, an agonist for FPR1-3 significantly reduces NFATc1 nuclear translocation, bone resorption and TRAP activity. The FPR family is essential for chemoattraction and immune responses (Migeotte et al. 2006) and are highly expressed in human monocytes (Hansen et al. 2024). Other work has shown that FPRs may have a role in bone cells. *Fpr1* knockout mice have reduced osteogenesis and bone fracture healing, and FPR1 activation promotes human osteoblast differentiation (Shin, et al. 2011; Yang, et al. 2024). Additionally, the FAM19A5 cytokine that activates FPR1 and FPR2 inhibits osteoclast formation and RANKL-induced gene expression in mouse cells (Park et al. 2017). Our studies indicated that the fMLF effects on human osteoclasts were mediated by FPR1 and could involve FPR3, while FPR2 antagonism had no effect on fMLF-mediated responses. This contrasts with previous studies in mice that indicate that the WRW4 antagonist prevents FAM19A5 effects on osteoclasts (Park et al. 2017). These differences could be due to differences in FPR gene expression, as mice express eight different FPRs compared to the three in humans (Gao, et al. 1998). Therefore, it may be difficult to extrapolate findings on specific FPRs between species. Alternatively, there may be ligand-specific differences as FPR1 has an ∼400-fold higher affinity for fMLF than FPR2 (Ye, et al. 1992). The role of FPR3 has not been investigated in bone cells and our studies with dual FPR1/2 antagonists proved too toxic to pursue in osteoclast activity assays. Further studies with other compounds or receptor-specific siRNA could elucidate whether other FPRs have a role in human osteoclasts.

In conclusion, these studies have demonstrated that HCI is a viable alternative to assess the effects of compounds on osteoclast activity. These assays could be utilised for high-throughput compound screening to rapidly assess novel mediators of osteoclast activity and identify compounds that may have utility in osteoporosis treatment.

## Author contribution statement

Conceptualization: CMG

Methodology: MLP, JC, CMG

Resources: AC, RH

Investigation: MLP, RAW, ZA, MH, CMG

Supervision: MF, CMG

Writing – original draft: MLP, CMG

Writing – review and editing: All authors

## Data and materials availability

All data needed to evaluate the conclusions in the paper are present in the paper and/or the Supplementary Materials. Raw data will be made available upon request. RNA-sequencing data generated from cultured primary osteoclasts derived from eight human donors have been deposited in the Gene Expression Omnibus (GEO) database under accession code GSE246769. Processed scRNA-seq data are available at the open science framework: https://osf.io/9xys4/. The analysis platform for quantifying nuclear fluorescence using arivis is available at the open science framework https://osf.io/43a8h/.

